# Proteolytic Activation of Human-specific Olduvai Domains by the Furin Protease

**DOI:** 10.1101/2021.07.06.450945

**Authors:** Ashley Pacheco, Aaron Issaian, Jonathan Davis, Nathan Anderson, Travis Nemkov, Beat Vögeli, Kirk Hansen, James M. Sikela

## Abstract

Olduvai protein domains (formerly DUF1220) show the greatest human-specific increase in copy number of any coding region in the genome and are highly correlated with human brain evolution and cognitive disease. The majority of human copies are found within four *NBPF* genes organized in a variable number of a tandemly arranged three-domain blocks called Olduvai triplets. Here we show that these human-specific Olduvai domains are posttranslationally processed by the furin protease, with a cleavage site occurring once at each triplet. These findings suggest that all expanded human-specific *NBPF* genes encode proproteins consisting of many independent Olduvai triplet proteins which are activated by furin processing. The exceptional correlation of Olduvai copy number and brain size taken together with our new furin data, indicates the ultimate target of selection was a rapid increase in dosage of autonomously functioning Olduvai triplet proteins, and that these proteins are the primary active agent underlying Olduvai’s role in human brain expansion.

## INTRODUCTION

Olduvai protein domains (formerly called DUF1220) are approximately 65 amino acids in length and are found almost exclusively within *NBPF* genes (Sikela and van Roy 2018). Studies have shown that Olduvai copy number is linearly associated with primate brain size, neuron number, and several other brain size-related phenotypes (Dumas and Sikela 2009; Dumas, et al. 2012; Keeney, et al. 2015; Keeney, et al. 2014; Zimmer and Montgomery 2015). In humans, Olduvai copy number is associated in a dosage-dependent manner to brain size, gray matter volume, and cognitive aptitude (Davis, et al. 2015b; Dumas, et al. 2012). Olduvai domains have undergone strong positive selection among primates and Olduvai copy number increases with evolutionary proximity to humans (Popesco, et al. 2006). Non-primate mammals have 1-9 copies, monkeys have 25-40 copies, great apes have 90-125 copies, and humans have 250-350 copies (O’Bleness, et al. 2012). The hyper-amplification of Olduvai in humans represents the largest human-specific increase in copy number of any coding region in the genome (O’Bleness, et al. 2012; Popesco, et al. 2006; Zimmer and Montgomery 2015).

The majority of Olduvai domains follow a unique genomic organization that allows copy number to be increased both rapidly and in an open-ended manner (Keeney, et al. 2014; O’Bleness, et al. 2012). Olduvai domains are encoded by a small and large exon doublet (Finn, et al. 2014; Popesco, et al. 2006; Vandepoele, et al. 2005) and can be phylogenetically subdivided into six primary clades: conserved (CON1-3) and human lineage-specific (HLS1-3) (O’Bleness, et al. 2012). The typical order of domain subtypes within human *NBPF* genes is: one to several CON1 domains; one CON2 domain; one-to-many instances of a triplet composed of HLS1, HLS2, and HLS3 domains; lastly one CON3 domain. The three-domain block of HLS sequences is known as the Olduvai triplet (O’Bleness, et al. 2012; Sikela and Searles Quick 2018).

The rapid and extreme human lineage-specific Olduvai increase is predominantly the result of tandem additions of Olduvai triplets within *NBPF* genes supplemented by gene duplications after Olduvai triplet expansion (Fiddes, et al. 2019; O’Bleness, et al. 2012). The proposed mechanism for the human expansion of *NBPF* genes is through a recombination event between HLS1 and CON3 which introduced a putative G-quadraplex (pG4) sequence pattern (Heft, et al. 2020). G4 structures promote instability which in conjunction with selective pressure is thought to have contributed to the human-specific extreme expansion of *NBPF* genes through tandem additions of Olduvai triplets (Heft, et al. 2020). In *NBPF* genes that contain one or more Olduvai triplets, the first triplet contains a canonical HLS1 small exon (HLS1-C), while every additional triplet contains a larger HLS1 small exon (HLS1-E) that is also found in the CON3 subtype (Heft, et al. 2020). In addition to the pG4 sequence, the extended exon of HLS1 also contains additional protein processing elements that have importance for human specific Olduvai function and will be discussed in this work.

The Olduvai protein domain family has been proposed to represent a genomic trade-off where selective pressure for increased copies resulted in increased brain size and cognition but, in certain situations, copies can also vary in deleterious ways (Sikela and Searles Quick 2018). Indeed, Olduvai copy number variation has been linked with multiple brain disorders including autism spectrum disorder (ASD), macrocephaly, microcephaly and schizophrenia (Davis, et al. 2019; Davis, et al. 2015a; Davis, et al. 2014; Dumas and Sikela 2009; Dumas, et al. 2012; Searles Quick, et al. 2015). Genomic data indicates that there are 23 human *NBPF* genes that carry from 5 to 60 tandemly arranged Olduvai domains. Such an arrangement suggests that the proteins encoded by human *NPBF* genes would span a wide range of sizes. Surprisingly, this is not the case. Rather Western analysis shows only a single primary band of ~36 kDa. This result has remained an enigma since it was first observed approximately 15 years ago (Popesco, et al. 2006). Here we present novel furin processing data confirming that expanded human-specific *NBPF* genes are posttranslationally cleaved, converting larger NBPF proproteins into smaller active protein units.

## RESULTS

### Mass Spec Analysis and Validation of Olduvai HLS Peptide Size

As mentioned above, predicted protein sizes of human NBPF proteins span a wide range, but western blot analysis of human tissues shows only a single band for the HLS domains at ~36kDa (Supplemental Figure 1A depicts previously published data from Popesco et al. 2006) (Popesco, et al. 2006). This product could contain 3-4 Olduvai domains and suggests that the functional size of HLS protein is smaller (Keeney, et al. 2014).

To further investigate this observation, the 36kDa band was enriched and analyzed by mass spectrometry-based proteomics. Chromatographic pre-enrichment of the 36kDa band combined with data-dependent global proteomic analysis identified a characteristic peptide of the Olduvai HLS subtype (Supplemental Figure 1B). Subsequent targeted proteomics analysis was able to identify additional peptides specific to NBPF including HLS domains (Supplemental Figure 2).

### Predicted Furin Processing Sites

Analysis of potential protease cut sites determined that translated HLS1-E exons contain a predicted furin cleavage site (Figure 1A). The HLS1-E exon, including the predicted furin site, is found within every expanded Olduvai triplet and is absent from all unexpanded triplets (i.e. those containing the HLS-C exon)(Figure 1B).

**Fig. 1.**
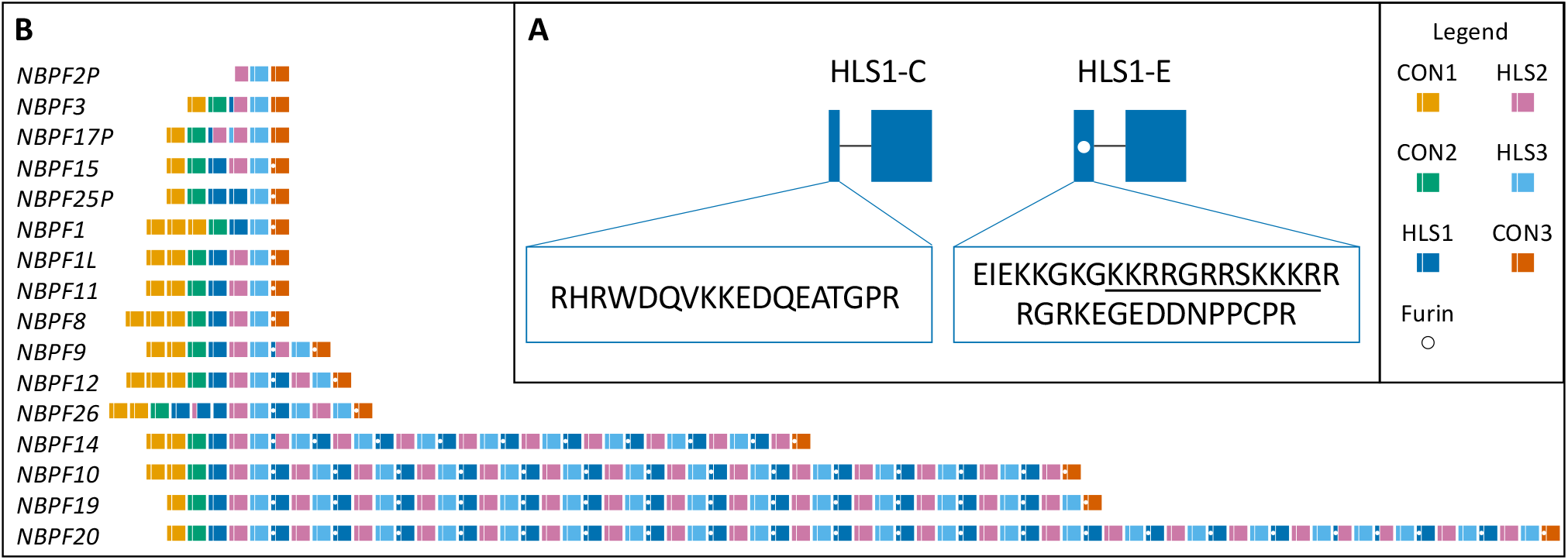
**(A)** Analysis of protease cleavage sites predicted a furin site located in the small exon of HLS1-E indicated by underlined amino acid sequence and the white dot in the figure. The small exons of canonical HLS1 (HLS1-C) and expanded HLS1 (HLS1-E) domains are enlarged to show the translated amino acid sequence. **(B)** All expanded NBPF Olduvai triplets contain a predicted furin cleavage site located within the small exons of HLS1-E and CON3 indicated by the white dots in the figure. Figure adapted from Heft et al., 2020.

### Furin Protease Processing Cleaves Extended HLS Domains

To evaluate potential furin processing of Olduvai triplets, both HLS1-C and HLS1-E were co-expressed with furin. A single band is observed for HLS1-C while a multiple banding pattern is observed for HLS1-E with furin co-expression (Figure 2A). These results confirm that extended HLS1 domains are proteolytically processed by furin from longer ‘proproteins’. The resulting predicted protein products from furin processing would include the longer N-terminal region of the protein including CON1 and CON2 and Olduvai triplets with HLS1-C, all Olduvai triplets containing the alternative HLS1-E, and finally the CON3 domain (Figure 2B). Furin processing of HLS1 would occur once at every Olduvai triplet within expanded *NBPF* genes generating numerous Olduvai triplet products from each gene. Confirmation of processing by furin resolves the long-standing question as to why only one protein band size is observed for HLS domains, despite the wide range of predicted protein sizes.

**Fig 2.**
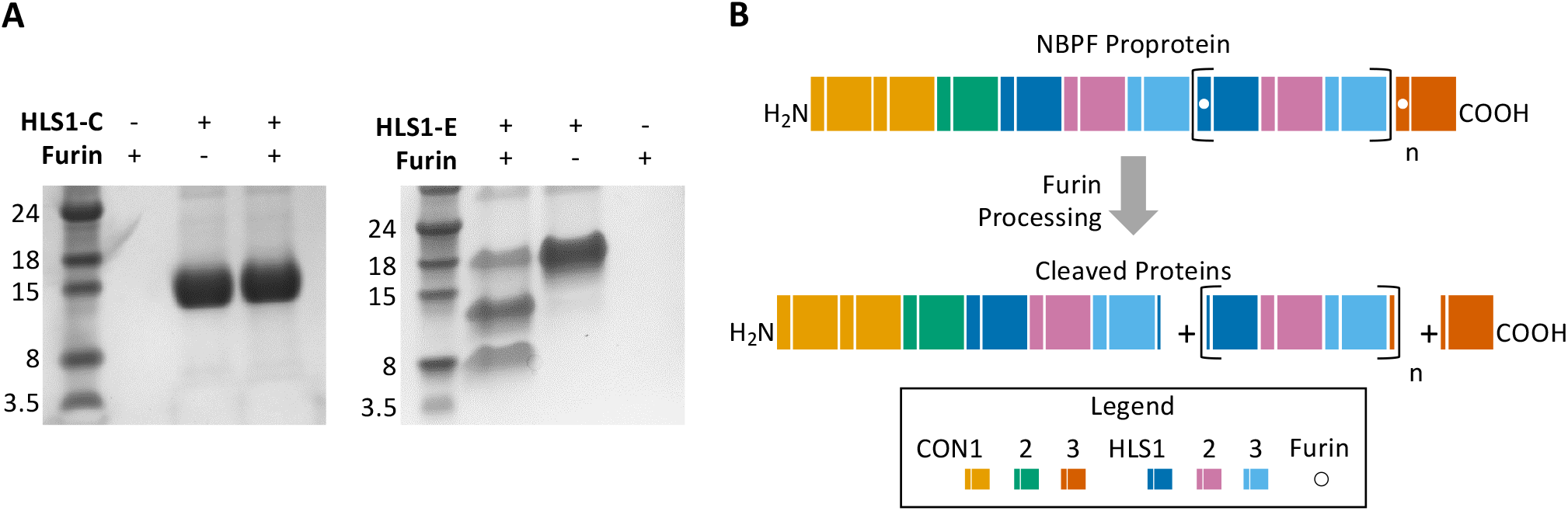
**(A)** Furin is shown to cleave expanded HLS1 (HLS1-E). To analyze furin cleavage of HLS1, both HLS1-C and HLS1-E were independently co-expressed with furin. A single banding pattern is shown when HLS1-C is co-expressed with furin (as seen on the left), whereas a multiple banding pattern is seen when HLS1-E is co-expressed with furin (as seen on the right). This finding confirms the previously predicted furin cleavage site that is located within HLS1-E. **(B)** Furin processing of the NBPF proprotein would result in three categories of peptides: the NBPF N-terminal peptide, “n” number of independent Olduvai triplet peptides, and the C-terminal CON3 peptide. A hypothetical NBPF protein with “n” number of Olduvai triplets is shown before and after furin processing. Furin cleavage sites are indicated by the white dots in HLS1-E and CON3.

The resulting predicted size for the expanded Olduvai triplet protein is approximately ~28kDa, which is lower than the observed ~36kDa band. While a higher than expected observed molecular weight (Mw) for this product could simply be due to structural peculiarities of the triplet protein, it may also indicate presence of posttranslational modifications (PTMs). It has been previously determined that Olduvai domains are intrinsically disordered (Issaian, et al. 2019; Wu, et al. 2020). As protein phosphorylation is the most common form of PTM, and intrinsically disordered proteins are disproportionately phosphorylated relative to other proteins (Bah and Forman-Kay 2016), predictive analysis for protein phosphorylation was conducted. Prediction of protein phosphorylation sites shows the presence of multiple predicted sites within the cleaved protein product, which could account for the increased observed Mw of the cleaved Olduvai triplet protein (Supplemental Figure 3). This finding is preliminary and further investigation will be needed to determine whether other forms of PTM are involved.

## DISCUSSION

The evolutionary hyper-amplification of the Olduvai protein domain family in humans represents one of the most dramatic human lineage-specific genomic events. Due to several recent studies, we are beginning to understand the specific steps of how this occurred primarily through the following two processes. The first is that human Olduvai expansion has employed a very rapid and highly expandable mechanism for increasing copy number by addition of tandemly arrayed Olduvai triplets (Heft, et al. 2020). The second being the addition of a furin cleavage site within each triplet which allows expanded copies to be readily converted via posttranslational processing to smaller independent active protein units, i.e., Olduvai triplets. It is likely that these events were driven by strong evolutionary selection pressures as they are seen within several human *NBPF* genes, and have added over 50 Olduvai triplets to the human genome over the past 2-3 million years (Fiddes et al 2019). Taken together, these findings indicate that, rather than simply raising copy number, the primary target of selection in humans has been to rapidly increase the dosage of autonomously functioning Olduvai triplet proteins, and these are the primary active agents of Olduvai function in humans.

Notably, processing by furin indicates that the N-terminal region including CON1 and CON2 as well as the C-terminal CON3 may be separated from extended HLS domains after translation. This finding suggests that CON and HLS domains may have independent functionality, which is supported by differing associations for various Olduvai domain subtypes. For example, one of the most striking and well-supported findings is the linear association between CON1 copy number and ASD severity (Davis, et al. 2019; Davis, et al. 2015a; Davis, et al. 2014). This link is not found with any other Olduvai subtypes, despite their being adjacent to CON1 in the NBPF proprotein. Independent function of the NBPF N-terminal protein product may have relevance to associations between CON1 and ASD severity and is to be discussed in a subsequent paper.

Finally, the new insights presented here resolve a long-standing question regarding the function of Olduvai domains in humans, and show how selection has favored increasing levels of individual Olduvai triplet proteins. How this dosage phenomenon is tied to the evolutionary increase in human brain size is an interesting future area of investigation.

## MATERIALS AND METHODS

### Mass Spec Analysis

Untargeted proteomic identification was performed on protein digests using liquid chromatography-tandem mass spectrometry as previously reported (Dzieciatkowska, et al. 2014). Briefly, nanoflow reverse-phase LC-MS/MS was performed using an Eksigent nanoLC2D system coupled to a LTQ Orbitrap-Velos mass spectrometer (Thermo Fisher). Data acquisition was performed using XcaliburTM (Version 2.1, Thermo Fisher) software. Raw data files were converted into Mascot generic files using PAVA. These peak lists were searched against SwissProt human database using Mascot server (Version 2.2.06, Matrix Science). Mass tolerances were 15 ppm for MS peaks, and 0.5 Da for MS/MS fragment ions.

### Enrichment of 36kDa Band

To enrich the 36kDa protein with antibody positivity for proteomics analysis, 100 mg of human brain tissue (Keeney, et al. 2015) was extracted and subjected to a series of fractionations during which the presence of the 36kDa band was followed by western analysis using an antibody directed against the HLS subtype sequence. Antibody positive fractions were then dissolved in 0.1% formic acid and analyzed by targeted proteomics. Multiple reaction monitoring MRM was performed using an LC-MS/MS system interfaced with a an UPLC system (QTRAP 5500 and Ultimate 3000, respectively, Sciex and Thermo Fisher Scientific). The mass spectrometer was run in positive ionization mode with the following settings: a source temperature of 200°C, spray voltage of 5300 V, curtain gas of 20 psi, and a source gas of 35 psi (nitrogen gas). Multiple SRM transitions were monitored using unit resolution in both Q1 and Q3 quadrupoles to maximize specificity. SRM assay optimization was performed with the aid of computer software (Skyline, Version 3.1). Collision energies and declustering potential were optimized for each transition. Method building and acquisition were performed using the instrument-supplied software (Analyst, Version 1.5.2, AB Sciex). Raw SRM data files were imported to Skyline Version 3.1 software for data processing.

### Protein Expression and Purification

Codon optimized gene fragments of HLS-C and HLS-E were cloned into pET21b(+) with the NdeI and XhoI restriction sites. Each expression construct contained an N-terminal MBP tag, a TEV cleavage site, and C-terminal 6xHis-tag. The plasmids were transformed and expressed in *Escherichia coli* strain Rosetta™ 2(DE3)pLysS cells with appropriate antibiotic selection. All bacterial expressed recombinant proteins were grown in Luria–Bertani (LB) broth.

HLS-C and HLS-E were expressed as follows. Three colonies from an overnight culture plate were resuspended in 1.5 L of LB with 0.001% Antifoam 204 and shaken at 26 °C for 14 hours. The A600 was measured to ensure the culture was below 0.5. Fresh LB was inoculated with the overnight culture (3:1) and shaken at 37 °C. The incubator temperature was lowered to 25 °C once the A600 reached 0.4. The culture was induced with 0.5 mM Isopropyl β- d-1-thiogalactopyranoside (IPTG) once the A600 reached 0.6 and shaken for 6 hours at 25 °C. Cells were harvested by centrifugation at 4 °C for 10 min at 4500 rpm.

HLS-C and HLS-E were purified as follows. Cell pellets were resuspended in 50 mM Tris (pH 7.5), 300 mM NaCl, 8 M urea, 2% glycerol, 10 mM imidazole, and 10 mM βME. Cell disruption was performed by sonication (7 x 20 s) at 4 °C. The lysates were clarified by centrifugation at 13,000 rpm for 30 min at 4 °C and loaded onto a pre-equilibrated column packed with Ni Sepharose™ excel resin. The column was washed with 5 CV of resuspension buffer and His-tagged proteins were eluted with 50 mM Tris (pH 7.5), 50 mM NaCl, 8 M urea, 3 % glycerol, and 400 mM imidazole. The protein sample was diluted and loaded onto a pre-equilibrated column packed with Source™ 15Q. Proteins were separated using the following gradient: 0 – 50% B (0 – 12 CV), 50 – 75 % B (12 – 15 CV), 75 – 100 % B (15 – 15.5 CV), 100 % B (15.5 – 16.5 CV) with buffer A (20 mM Tris (pH 8.0), 8 M urea) and buffer B (20 mM Tris (pH 8.0), 8 M urea, 1 M NaCl). The relevant fractions were pooled and dialyzed overnight at 4 °C against 20 mM Tris (pH 7.5), 50 mM NaCl, and 5 mM DTT. TEV protease (in-house) was added to the sample and incubated overnight at 4 °C with gentle rocking. The protein sample was loaded onto a pre-equilibrated column packed with Source™ 15Q. Proteins were separated using the following gradient: 0 – 50 % B (0 – 12 CV), 50 – 75 % B (12 – 15 CV), 75 – 100 % B (15 – 15.5 CV), 100 % B (15.5 – 16.5 CV) with buffer A (20 mM Tris (pH 8.0), 8 M urea) and buffer B (20 mM Tris (pH 8.0), 8 M urea, 1 M NaCl). Relevant fractions were pooled and dialyzed overnight at 4 °C against 20 mM HEPES (pH 7.5), 200 mM NaCl, and 5 mM DTT. The samples were concentrated to 1.5 mL using a 3000 MWCO concentrator (Sartorius) and diluted to 3 mL with urea buffer (20 mM Tris (pH 7.5), 300 mM NaCl, 8 M urea, 5 mM DTT). The samples were injected onto a Superose™ 6 Increase column (Cytiva) pre-equilibrated with native storage buffer (20 mM HEPES (pH 7.5), 200 mM NaCl, 5 mM DTT). Fractions containing pure target protein were pooled, concentrated, and stored at −80 °C.

### In-Solution Enzymatic Digestion

Both HLS-C and HLS-E protein at 1 mg/mL were incubated with human Furin (EC 3.4.21.75, P09958, obtained from NEB P8077) in digestion buffer (10 mM HEPES (pH 7.5), 0.2 mM CaCl2, 0.2 mM βME, 0.1% Triton X-100) for 16 hours at RT (25°C). Additional samples lacking either Olduvai Triplet or Furin were assembled as experimental controls. Samples were mixed with 1X LDS Sample Buffer (GenScript) supplemented with 100 mM βME and incubated at 95 °C for 5 min. Equal volumes of each sample were loaded onto a 4-12% SDS-PAGE gel along with the BLUEstainTM 2 (GoldBio) protein ladder. The gel was stained with SimplyBlueTM Safe Stain (Thermo Fisher Scientific) and visualized with a CCD equipped imager (Bio-Rad).

### *In-situ* Analyses

Furin site prediction was carried out by submitting full length NBPF protein sequences to the ProP – 1.0 server (Duckert, et al. 2004). Protein phosphorylation site prediction was conducted by submitting Olduvai triplet protein sequences containing cleaved HLS1-E exons to the NetPhos – 3.1 server(Blom, et al. 1999).

## ACKNOWLEDGEMENTS

We thank Ilea Heft, Morkos Henen and Natasia Paukovich for helpful discussions and Monika Dzieciatkowska for assistance with mass spectrometry experiments. This work was supported by the National Institute of General Medical Sciences at the National Institutes of Health (R01GM130694-01A1 to B.V.), and the National Institute of Mental Health at the National Institute of Health (R01MH108684 to J.M.S.).

## DATA AVAILABILITY STATEMENT

The data underlying this article are available in the article and in its online supplementary material.

## References

Bah A, Forman-Kay JD 2016. Modulation of Intrinsically Disordered Protein Function by Post-translational Modifications. J Biol Chem 291: 6696–6705. doi: 10.1074/jbc.R115.695056

Blom N, Gammeltoft S, Brunak S 1999. Sequence and structure-based prediction of eukaryotic protein phosphorylation sites. J Mol Biol 294: 1351–1362. doi: 10.1006/jmbi.1999.3310

Davis JM, Heft I, Scherer SW, Sikela JM 2019. A Third Linear Association Between Olduvai (DUF1220) Copy Number and Severity of the Classic Symptoms of Inherited Autism. Am J Psychiatry 176: 643–650. doi: 10.1176/appi.ajp.2018.18080993

Davis JM, Searles Quick VB, Sikela JM 2015a. Replicated linear association between DUF1220 copy number and severity of social impairment in autism. Hum Genet 134: 569–575. doi: 10.1007/s00439-015-1537-6

Davis JM, et al. 2014. DUF1220 dosage is linearly associated with increasing severity of the three primary symptoms of autism. PLoS Genet 10: e1004241. doi: 10.1371/journal.pgen.1004241

Davis JM, et al. 2015b. DUF1220 copy number is linearly associated with increased cognitive function as measured by total IQ and mathematical aptitude scores. Hum Genet 134: 67–75. doi: 10.1007/s00439-014-1489-2

Duckert P, Brunak S, Blom N 2004. Prediction of proprotein convertase cleavage sites. Protein Eng Des Sel 17: 107–112. doi: 10.1093/protein/gzh013

Dumas L, Sikela JM 2009. DUF1220 domains, cognitive disease, and human brain evolution. Cold Spring Harb Symp Quant Biol 74: 375–382. doi: 10.1101/sqb.2009.74.025

Dumas LJ, et al. 2012. DUF1220-domain copy number implicated in human brain-size pathology and evolution. Am J Hum Genet 91: 444–454. doi: 10.1016/j.ajhg.2012.07.016

Dzieciatkowska M, Hill R, Hansen KC. 2014. GeLC-MS/MS Analysis of Complex Protein Mixtures. In: Martins-de-Souza D, editor. Shotgun Proteomics: Methods and Protocols. New York, NY: Springer New York. p. 53–66.

Fiddes IT, Pollen AA, Davis JM, Sikela JM 2019. Paired involvement of human-specific Olduvai domains and NOTCH2NL genes in human brain evolution. Hum Genet 138: 715–721. doi: 10.1007/s00439-019-02018-4

Finn RD, et al. 2014. Pfam: the protein families database. Nucleic Acids Res 42: D222–230. doi: 10.1093/nar/gkt1223

Heft IE, et al. 2020. The Driver of Extreme Human-Specific Olduvai Repeat Expansion Remains Highly Active in the Human Genome. Genetics 214: 179–191. doi: 10.1534/genetics.119.302782

Issaian A, et al. 2019. Solution NMR backbone assignment reveals interaction-free tumbling of human lineage-specific Olduvai protein domains. Biomol NMR Assign 13: 339–343. doi: 10.1007/s12104-019-09902-0

Keeney JG, et al. 2015. DUF1220 protein domains drive proliferation in human neural stem cells and are associated with increased cortical volume in anthropoid primates. Brain Struct Funct 220: 3053–3060. doi: 10.1007/s00429-014-0814-9

Keeney JG, Dumas L, Sikela JM 2014. The case for DUF1220 domain dosage as a primary contributor to anthropoid brain expansion. Front Hum Neurosci 8: 427. doi: 10.3389/fnhum.2014.00427

O’Bleness MS, et al. 2012. Evolutionary history and genome organization of DUF1220 protein domains. G3 (Bethesda) 2: 977–986. doi: 10.1534/g3.112.003061

Popesco MC, et al. 2006. Human lineage-specific amplification, selection, and neuronal expression of DUF1220 domains. Science 313: 1304–1307. doi: 10.1126/science.1127980

Searles Quick VB, Davis JM, Olincy A, Sikela JM 2015. DUF1220 copy number is associated with schizophrenia risk and severity: implications for understanding autism and schizophrenia as related diseases. Transl Psychiatry 5: e697. doi: 10.1038/tp.2015.192

Sikela J, van Roy F 2018. Changing the name of the NBPF/DUF1220 domain to the Olduvai domain [version 2; peer review: 3 approved]. F1000Research 6. doi: 10.12688/f1000research.13586.2

Sikela JM, Searles Quick VB 2018. Genomic trade-offs: are autism and schizophrenia the steep price of the human brain? Hum Genet 137: 1–13. doi: 10.1007/s00439-017-1865-9

Vandepoele K, Van Roy N, Staes K, Speleman F, van Roy F 2005. A novel gene family NBPF: intricate structure generated by gene duplications during primate evolution. Mol Biol Evol 22: 2265–2274. doi: 10.1093/molbev/msi222

Wu H, Zhai LT, Guo XX, Rety S, Xi XG 2020. The N-terminal of NBPF15 causes multiple types of aggregates and mediates phase transition. Biochem J 477: 445–458. doi: 10.1042/BCJ20190566

Zimmer F, Montgomery SH 2015. Phylogenetic Analysis Supports a Link between DUF1220 Domain Number and Primate Brain Expansion. Genome Biol Evol 7: 2083–2088. doi: 10.1093/gbe/evv122

