## Supplemental Figures for "Proteolytic Activation of Human-specific Olduvai Domains by the Furin Protease"

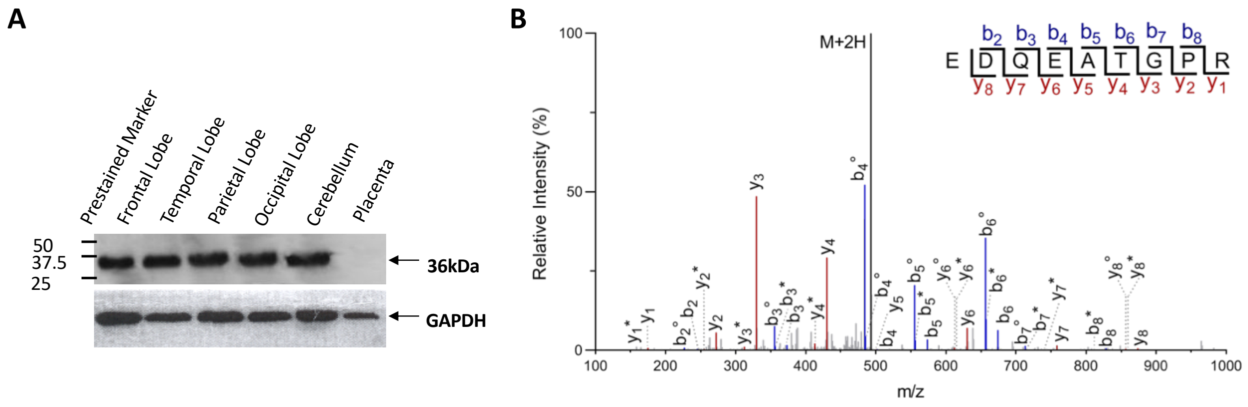


**Supplemental Figure 1. (A)** Data was previously published in Popesco et al., 2006 (Figure 4B). Western blot analysis using an antibody against a primate-specific Olduvai (formerly DUF1220) domain in human brain tissue shows a heavy band at ~36kDa. **(B)** Spectrum assignment for Olduvai HLS-containing peptide identified from fractionated human brain tissue. For C-terminal y ion (red) and N-terminal b ion (blue) peptide fragments derived from the in-tact peptide (M+2H, black) the C-terminal, * denotes the loss of $NH_3$, and ° denotes the loss of $H_2O$.

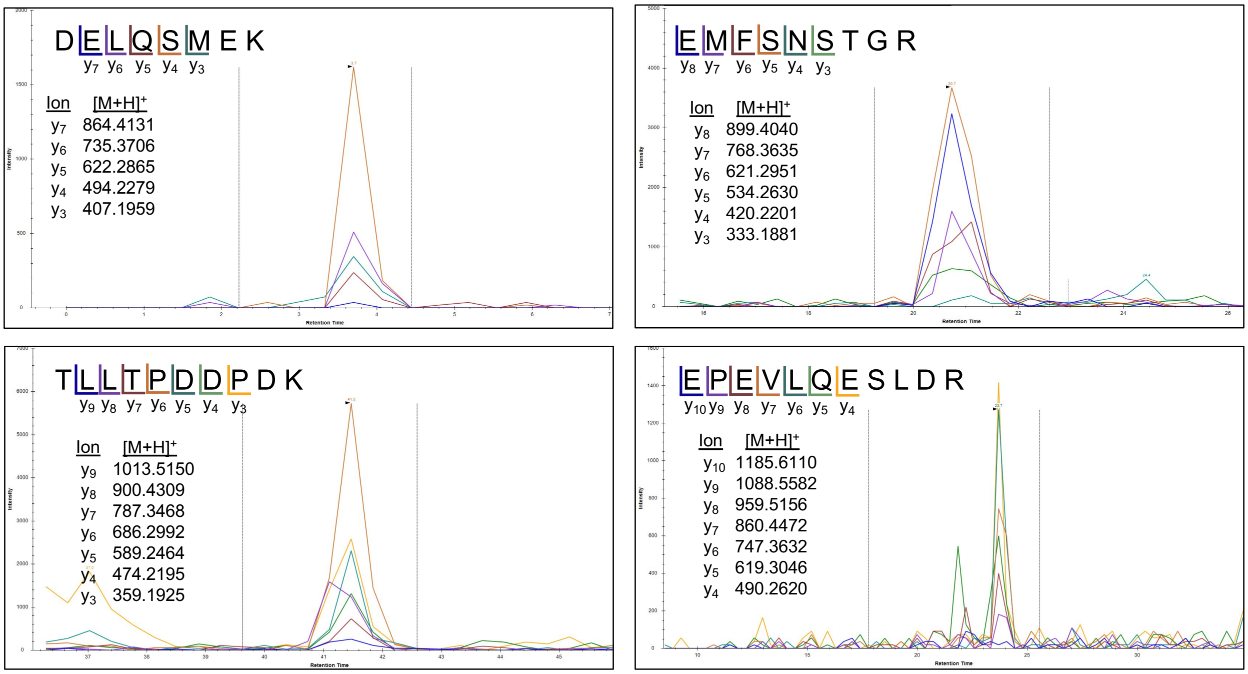


**Supplemental Figure 2.** Results depict additional NBPF peptides identified by targeted proteomics analysis within human brain tissue enriched for the 36kDa band confirming that the 36kDa band contains Olduvai sequences.

NPBF14 Olduvai Predicted Phosphorylation Sites

SKKERRRGRKEGEEDQNPPCPRLSRELLDEKGPEVLQDSLDRCYSTPSGCLELTDSCQPY
RSAFYVLEQQRVGLAVDMDEIEKYQEVEEDQDPSCPRLSRELLDEKEPEVLQDSLDRCY
STPSGYLELPDLGQPYSSAVYSLEEQYLGLALDVDRIKKDEEEEEDQDPPCPRLSRELLE
VVEPEVLQDSLDRCYSTPSSCLEQPDSCQPYGSSFYALEEKHVGFSLDVGEIEKKGKGKK
RRGRR

Key

HLS1 HLS2 HLS3

Phosphorylation

**Supplemental Figure 3.** Furin cleaved NBPF14 HLS domains are depicted. Yellow highlights show locations of predicted phosphorylation sites from NetPhos.
